# Microclimate and larval habitat density predict adult *Aedes albopictus* abundance in urban areas

**DOI:** 10.1101/615625

**Authors:** Michelle V. Evans, Carl W. Hintz, Lindsey Jones, Justine Shiau, Nicole Solano, John M. Drake, Courtney C. Murdock

## Abstract

The Asian tiger mosquito, *Aedes albopictus*, transmits several arboviruses of public health importance, including chikungunya and Zika. Since its introduction to the United States in 1985, the species has invaded over forty states, including temperate areas not previously at risk of *Aedes*-transmitted arboviruses. Mathematical models incorporate climatic variables in predictions of site-specific *Ae. albopictus* abundances to identify human populations at risk of disease. However, these models rely on coarse resolutions of environmental data that may not accurately represent the climatic profile experienced by mosquitoes in the field, particularly in climatically-heterogeneous urban areas. In this study, we pair field surveys of larval and adult *Ae. albopictus* mosquitoes with site-specific microclimate data across a range of land use types to investigate the relationships between microclimate, density of larval habitat, and adult mosquito abundance and determine whether these relationships change across an urban gradient. We find no evidence for a difference in larval habitat density or adult abundance between rural, suburban, and urban land classes. Adult abundance increases with increasing larval habitat density, which itself is dependent on microclimate. Adult abundance is strongly explained by microclimate variables, demonstrating that theoretically derived, lab-parameterized relationships in ectotherm physiology apply to the field. Our results provide support for the continued use of temperature-dependent models to predict *Ae. albopictus* abundance in urban areas.

## INTRODUCTION

The Asian tiger mosquito, *Aedes albopictus*, is an invasive mosquito that became established in the United States following its introduction in 1985.^1,2^ *Ae. albopictus* can transmit several pathogens of public health importance, including La Crosse,^3^ dengue,^4,5^ and chikungunya viruses.^6^ Unlike another vector of these diseases, *Aedes aegypti*, which originated in East Africa, *Ae. albopictus* originated from a temperate area of Asia and is able to survive in cooler climates than *Ae. aegypti.* Following initial establishment in Texas, *Ae. albopictus* has invaded over 40 states,^7^ and models predict its range will expand as the climate warms.^8,9^ At present, established populations of *Ae. albopictus* are found in the United States as far north as Connecticut and New York,^10,11^ well outside the present range of *Ae. aegypti*. *Ae. albopictus* is implicated in transmission cycles of dengue and chikungunya in the Mediterranean region of Europe,^12,13^ which suggests that temperate regions of the US may be similarly vulnerable.

Given the potential role of *Ae. albopictus* in disease transmission, it is important to understand what factors influence its abundance. *Ae. albopictus* is sensitive to variation in temperature due to temperature-dependent life history traits such as development rates, fecundity, and survival.^14–16^ Climate or meteorological predictors are widely used in mechanistic models and statistical models.^17–22^ Models leverage these relationships to predict mosquito presence, population growth rates, and abundances based on temperature metrics derived from weather stations or remotely-sensed datasets. However, urban landscapes are composed of a variety of land classes (e.g. residential, developed, vegetated), which vary in their microclimates at fine spatial scales less than 1 × 1 km.^23–25^ This difference in microclimate can alter mosquito population growth rates,^26,27^ leading to variation in population abundances that may be missed by models that rely on coarser spatial data.

Additionally, adult abundance may be determined by the abundance of larval habitat. *Ae. albopictus* is fairly non-discriminate in its habitat use and larvae are found in both natural and artificial containers.^11,28,29^ Several studies have found that adult abundance is positively related to the availability of larval habitats.^30,31^ This relationship is also the basis for larval source reduction techniques widely used in vector control.^32^ Urban microclimates can co-vary with mosquito larval habitat density, which may differ in quality and quantity across urban land-use.^26,30^ Thus, when studied independently, the relative roles of microclimate and larval habitat may be confounded.

Here, we combine field surveys of larval habitat and adult mosquito abundances with microclimate data to investigate how microclimate and the availability of larval habitat contribute to changes in adult *Ae. albopictus* abundance across an urban landscape. We aim to answer the following questions:

1. Does the density of larval habitat positive for *Ae. albopictus* change across urban land classes?
2. Does the abundance of *Ae. albopictus* adults change across urban land classes?
3. What is the relationship between microclimate and adult abundance?
4. What is the relationship between larval habitat and adult abundance?

By investigating these relationships, our results inform if and how predictive models should include microclimate variables and data on larval habitat from the field in their predictions of adult *Ae. albopictus* abundance. Further, these results can help determine whether variation in land class alters the spatial distribution of *Ae. albopictus*, and if this omitting this fine-scale variation may be leading to bias in models.

## MATERIALS & METHODS

The study was conducted between June 2016 and December 2017 in Athens-Clarke County, GA, USA. Athens-Clarke County is an urbanized area in a matrix of rural forested and agricultural land, representing a wide range of land classes. Following previous work,^33^ we used an impervious surface map (NLCD) to select three replicate 30 × 30 m sites each of low (0-5%), intermediate (6 - 40%), and high (41-100%) impervious surface (Fig. 1). Percent impervious surface, an accurate predictor of land surface temperature,^34^ was chosen to ensure the sites exhibited the full range of microclimates present in the city.

**Figure 1.**
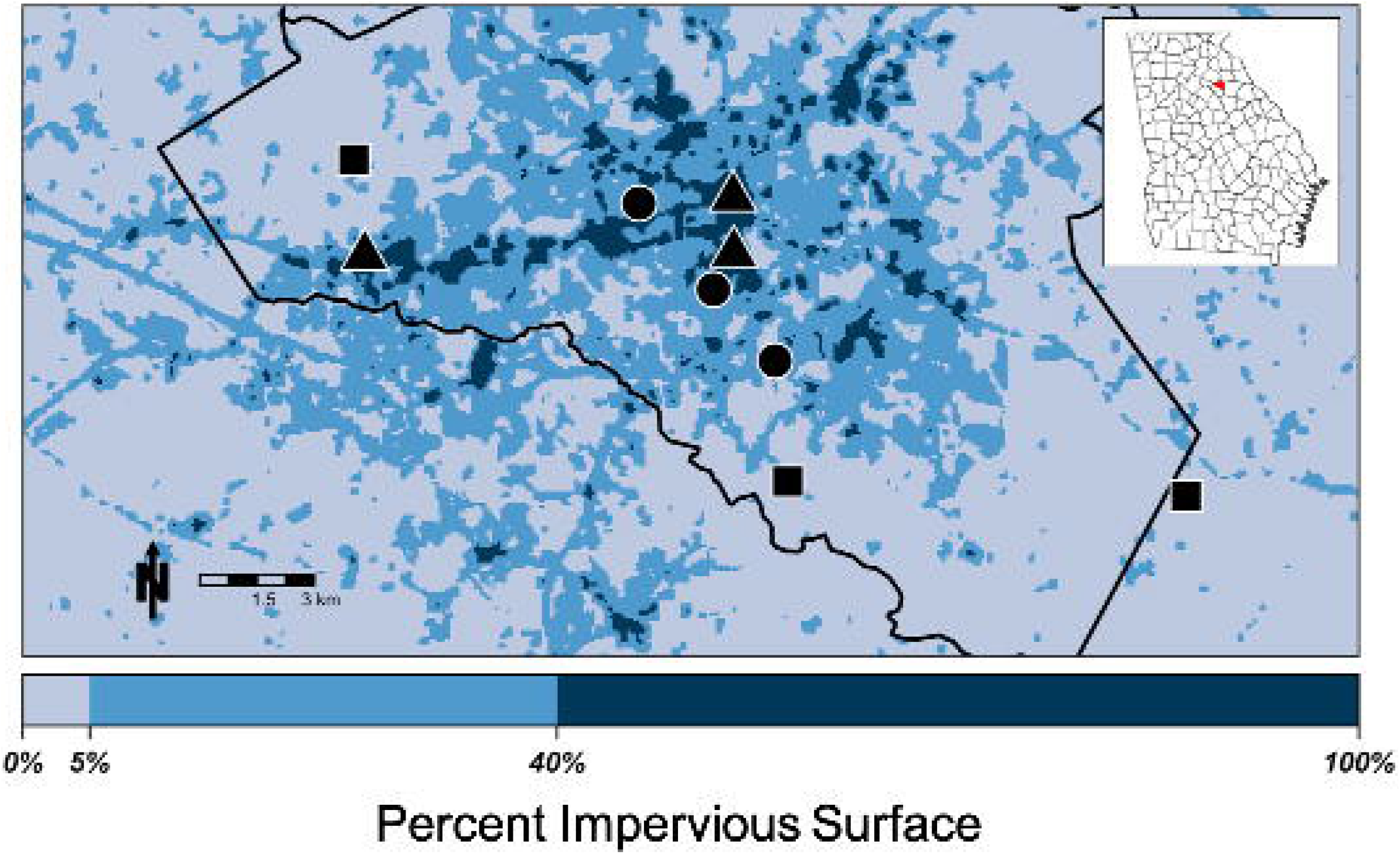
Map of sites in Athens, GA. Symbols represent land classes (square: rural, circle: suburban, triangle: urban). Color shading represents the amount of impervious surface within the 210 m focal area of each pixel, as illustrated on the color bar on the bottom. Athens-Clarke County is outlined in black and its location within the state is shown in the inset map of Georgia.

### Aquatic Immature Surveys

At each site, we conducted surveys of the aquatic immature stages (larvae, pupae) bi-weekly from June - December 2016 and April - December 2017 to measure the density of positive larval habitats (e.g. number of larval habitats positive for *Ae. albopictus* larvae per 100m radius site). Each site was sampled within one day, and the full sampling period of surveys took place over several days, with a sampling period referring to the week in which surveys were conducted. Study areas were defined as a 100m radius surrounding the center of the focal 30 × 30m site. Each study area was inspected for the presence of standing water (i.e. puddles, ponds, artificial containers). Each body of water was assessed for the presence of immature mosquitoes visually and using dipping methods. If immature mosquitoes were present, samples were collected from that habitat. Because adult mosquitoes were sampled concurrently with larval habitat sampling, destructive sampling could bias the adult catch rate. For this reason, we collected measures of presence or absence of *Ae. albopictus* per habitat by sampling a subset of the immature mosquitoes per habitat (ranging from 5 - 27 individuals per habitat). Immature mosquitoes were kept separated by habitat and returned to the lab, where they were placed in 50 - 100 mL deionized water in 8 oz. glass jars (Ball) and provided fish food (Hikari Cichlid Gold Mini Pellet) *ad libitum* to ensure high emergence rates. Larvae and pupae were reared to adulthood in an incubator (Percival Scientific) at 27 ± 0.5 °C, 80 ± 5 % relative humidity, and a 12:12 h light:dark photocycle. Once mosquitoes emerged, they were immediately frozen at −20 °C, separated by sex, and identified to species following Darsie and Ward.^35^ A habitat was determined ‘positive’ for *Ae. albopictus* during a sampling period if a male or female *Ae. albopictus* mosquito was identified as emerging from the habitat.

### Adult Trapping

At each site, we trapped adults either bi-weekly (during the season of highest mosquito activity, June - Nov 2016 and June - Nov 2017) or monthly (Dec 2016 - May 2017 and Dec 2017). During the period of highest mosquito activity, adult trapping was conducted within one week of immature surveys. One BG Sentinel-2 (Biogents, Germany) mosquito trap was deployed in the center of each 30 × 30m site for two consecutive trap days per sampling period. Mosquito traps were baited with a BG-Lure cartridge (Biogents, Germany) and an octenol (1-Octen-3-ol) lure inside the trap. Trapping was not conducted during precipitation events, and traps were placed under the cover of vegetation to increase catch rates. Because *Ae. albopictus* is a day biting mosquito, the traps were run (with a battery powered fan) from the hours of 06:00h to 22:00h. After each trap day, catch bags were collected and replaced with a new catch bag to reduce destruction of samples. Collected adults were taken back to the laboratory, frozen in a −20 °C freezer, and separated by sex and identified to species following Darsie and Ward.^35^ Abundances for both trap days were combined to calculate the total abundance for that sampling period. The date of that sampling period is defined as the day on which the second catch bag was collected.

### Microclimate Variables

Within each 30 × 30m site, we evenly distributed six data loggers (Monarch Instruments, Amherst, NH, USA: Radio Frequency Identification (RFID) Temperature Track-It Logger) to measure microclimate (e.g. site-specific climatic variables). Data loggers were placed in full shade under vegetation, approximately 0.9 m above the ground. The loggers recorded instantaneous temperature and relative humidity at ten-minute intervals. From the ten-minute data, we calculated daily minimum, mean, and maximum values for both temperature and relative humidity for each logger. These values were then averaged across all six loggers for each site.

### Data Analyses

To determine if the density of positive *Ae. albopictus* larval habitat differed across land class, we used a generalized linear mixed model (GLMM) to test for the effect of land class on the density of positive larval habitats, including site as a random effect. The model included the week number of the study period as a basis-spline (B-spline) function to account for seasonal differences in mosquito catch rates. The B-spline function allows a curve to be fit using maximum likelihood without pre-specifying a function.^36^ A similar model was used to explore the effect of land class on *Ae. albopictus* adult abundance, again including site as a random effect and the week number of the study period as a B-spline function. Both models used a negative binomial distribution in which the variance increases quadratically with the mean:

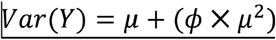

where μ is the mean and ϕ is the dispersion parameter of the distribution.^37^ Models used a logarithmic link function. The statistical significance of land class’ effect was assessed by comparing fitted models to a null model that did not include land class as a predictor variable using a likelihood ratio test.

We used univariate GLMM’s to investigate the effect of the microclimate variables on the density of positive larval habitat and adult abundance. Microclimate variables were fit using a B-spline function to allow for non-linear relationships and site was included as a random effect. All models were fit with the quadratic variance form of the negative binomial distribution described above and a logarithmic link function. We averaged each microclimate variable over the seven days prior to surveying to account for the fact that captured mosquitoes likely developed and emerged within that time period. We explored using two different lag widths, seven and fourteen days, in the models. Resulting models did not differ significantly, and so a lag of 7 days was used. This agrees with prior work in this system that found mosquito development rates to range from 7 - 10 days during periods of high mosquito activity.^27^ This resulted in the following variables: mean weekly temperature and relative humidity, minimum weekly temperature and relative humidity, and maximum weekly temperature. Maximum weekly relative humidity was excluded from the analysis because 226 out of 234 trap periods had a maximum value of 100% relative humidity. We included the day-of mean relative humidity value in models of adult abundance to control for mosquito activity on that trap day. We assessed the statistical significance of each microclimate variable by comparing fitted models to a null model that did not include the variable as a predictor variable using a likelihood ratio test.

We also tested for the effect of the density of positive *Ae. albopictus* larval habitat (the number of larval habitats that had *Ae. albopictus* larvae present per site) on adult abundance within a site and sampling period. A GLMM was fit including the density of positive habitats as a predictor variable and site as a random effect. We fit the model with the same negative binomial distribution and logarithmic link function as described above.

All GLMMs were fit using the *glmmTMB* package in R version 3.5.2.^38,39^ Scaled residuals of the models were inspected for overdispersion and uniformity using the *DHARMa* package.^40^ Code and data to reproduce analyses are deposited on the figshare repository (https://figshare.com/s/8ea9037a5f39bd4a4961).

## RESULTS

A total of 1107 adult female *Ae. albopictus* mosquitoes were sampled from May 2016 - December 2017, encompassing 468 trap nights over two seasons of mosquito activity. This resulted in 26 adult sampling events for each of the nine sites, or 78 sampling events per land class. We sampled each site for larval habitat a total of 21 times and found 217 habitats positive for *Ae. albopictus* across all nine sites.

### Land Class and Season

The density of larval habitats positive for *Ae. albopictus* was highly seasonal, peaking in June - August of both years (Fig. 2). The best fitting B-spline used a three-degree polynomial, and the effect of sampling week was significant (χ^2^ = 37.023, df = 3, p-value < 0.001). While suburban sites tended to have a higher density of positive larval habitat than rural and urban sites, this difference was not significant. A null model without land class as a predictor variable was not significantly different from the full model (χ^2^ = 4.34, df = 2, p-value = 0.110) and predictive performance was similar (R^2^_*NULL*_ = 0.503, R^2^_FULL_ = 0.483).

**Figure 2.**
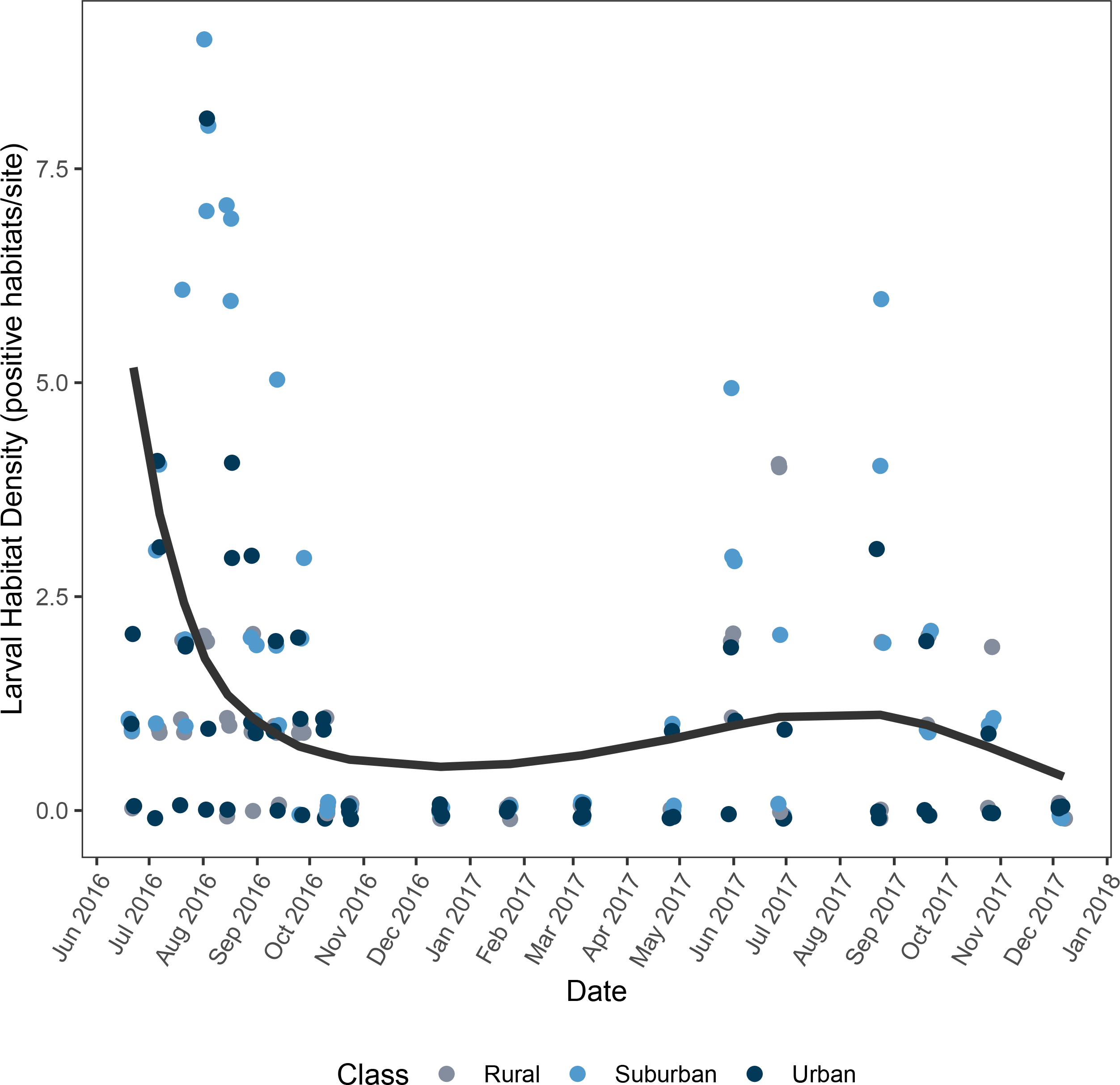
Positive larval habitat density across land class and time. Lines represent the fitted model averaged across all sites. Raw data are represented by the points, and randomly jittered to improve visibility.

We found evidence of very strong seasonality in adult *Ae. albopictus* density across all sites, with densities peaking in July and August of both years (Fig. 3). There was a significant effect of the sample week on adult density (χ^2^ = 112.050, df = 4, p-value < 0.001) and the best fitting B-spline had a four-degree polynomial. There was no evidence for a difference in adult *Ae. albopictus* density across land class. The null model without land class as a predictor variable was not significantly different from the full model (χ^2^ = 0.602, df = 2, p-value = 0.740) and performed similarly (R^2^_*NULL*_ = 0.813, R^2^_FULL_ = 0.813).

**Figure 3.**
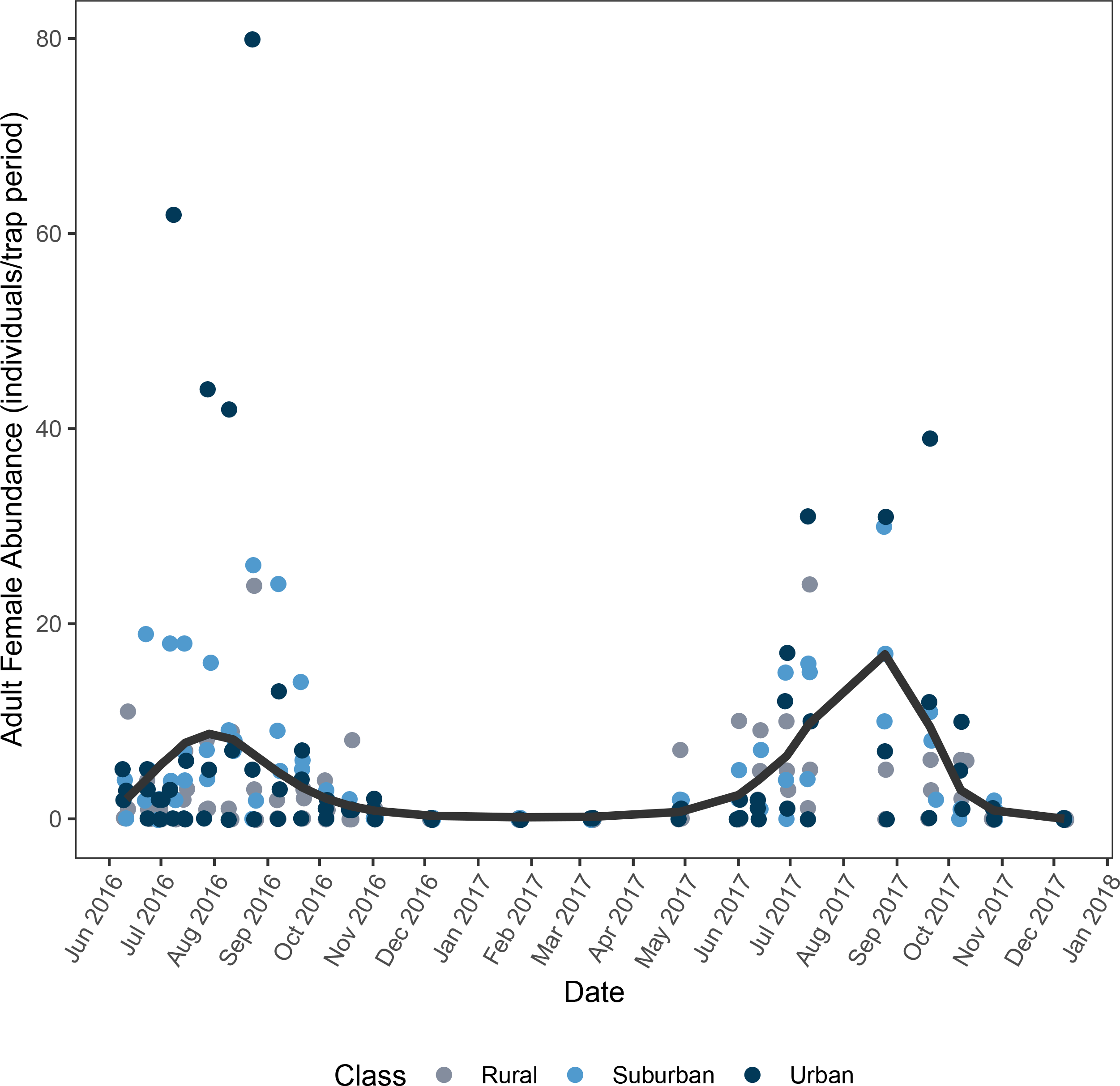
Adult female *Ae. albopictus* abundance across time. Lines represent the fitted model averaged across all sites. Raw data are represented by the points, and randomly jittered to improve visibility.

### Microclimate and Larval Habitat Density

Univariate analyses revealed a significant, non-linear relationship between all microclimate variables and the density of positive *Ae. albopictus* habitat (Table 1, Fig. 4). The density of positive larval habitat increased with increasing minimum, mean, and maximum temperatures (Fig. 4). Larval habitat increased with increasing minimum relative humidity until approximately 60% relative humidity, after which increasing relative humidity was associated with fewer larval habitats (Fig. 4). The relationship between mean relative humidity and the density of larval habitats was similarly unimodal, although its optimum neared 100% relative humidity. Importantly, the functional forms of these relationships differ from those between the microclimate variables and adult abundance. This difference suggests that the effects of microclimate on oviposition behavior and habitat availability differs from the effect of microclimate on mosquito emergence and adult longevity.

**Table 1.** Results of likelihood ratio tests comparing null model to univariate GLMMs containing microclimate variables to predict the density of *Ae. albopictus-*positive larval habitat. All microclimate variables were strong predictors of larval habitat density. GLMM was calculated across 9 sites (random effect), and within-site *n* = 21. We calculated conditional R^2^ following Schielzeth & Nakagawa (2013).^60^

| Variable | df | $\chi^2$ | p-value | Conditional $R^2$ |
| --- | --- | --- | --- | --- |
| Min. Temperature | 3 | 93.516 | <0.0001 | 0.764 |
| Mean Temperature | 3 | 92.189 | <0.0001 | 0.830 |
| Max. Temperature | 3 | 61.480 | <0.0001 | 0.855 |
| Min. Relative Humidity | 3 | 52.522 | <0.0001 | 0.695 |
| Mean Relative Humidity | 3 | 24.776 | <0.0001 | 0.506 |

**Figure 4.**
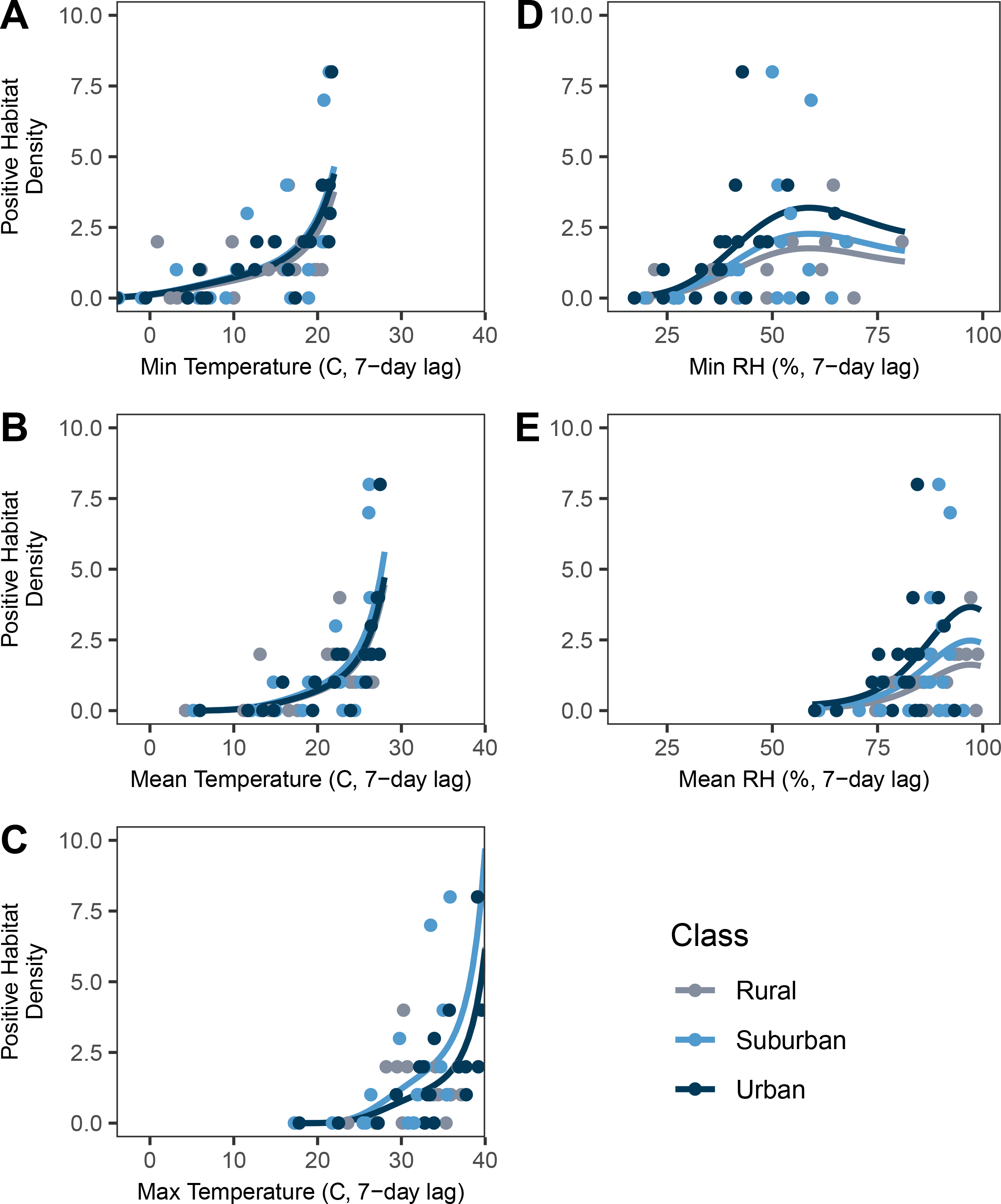
Functional relationship between microclimate variables and the density of positive *Ae. albopictus* habitat for a representative site of each land class. Functional relationships were the same across all land classes, and larval habitat density did not differ across land class. Daily minimum, mean, and maximum temperatures are on the left side (a,b,c) and daily minimum and mean relative humidity are on the right side (d,e). Lines represent fitted regression lines and raw data are represented by the circles. Because maximum relative humidity did not vary, no regression line was fitted.

### Microclimate and Adult Abundance

Univariate analyses revealed that all five microclimate variables significantly influenced adult abundances (Table 2). The relationships between microclimate variables and adult abundance were non-linear for all variables (Fig. 5). Mean and minimum relative humidity had a 3rd-order B-spline fit that increased exponentially as relative humidity approached 100% (Fig. 5). All three temperature variables also had a third-order B-spline fit, evidence of a non-linear relationship. Minimum daily temperature was similar to relative humidity in that it was an increasing function across the range measured in this experiment (−3.75 to 23.10°C). The functional relationships between mean and maximum temperature and adult abundance were unimodal, decreasing after an optimal temperature threshold was reached (Fig. 5).

**Table 2.** Results of likelihood ratio tests comparing null model to univariate GLMMs containing microclimate variables to predict adult female abundance. All microclimate variables were strong predictors of adult abundance. GLMM was calculated across 9 sites (random effect), and within-site *n* = 26. We calculated conditional R2 following Schielzeth & Nakagawa (2013).^60^

| Variable | df | $\chi^2$ | p-value | Conditional $R^2$ |
| --- | --- | --- | --- | --- |
| Min. Temperature | 3 | 104.27 | <0.001 | 0.835 |
| Mean Temperature | 3 | 110.94 | <0.0001 | 0.847 |
| Max. Temperature | 3 | 96.50 | <0.0001 | 0.910 |
| Min. Relative Humidity | 3 | 49.79 | <0.0001 | 0.608 |
| Mean Relative Humidity | 3 | 16.257 | 0.001 | 0.530 |

**Figure 5.**
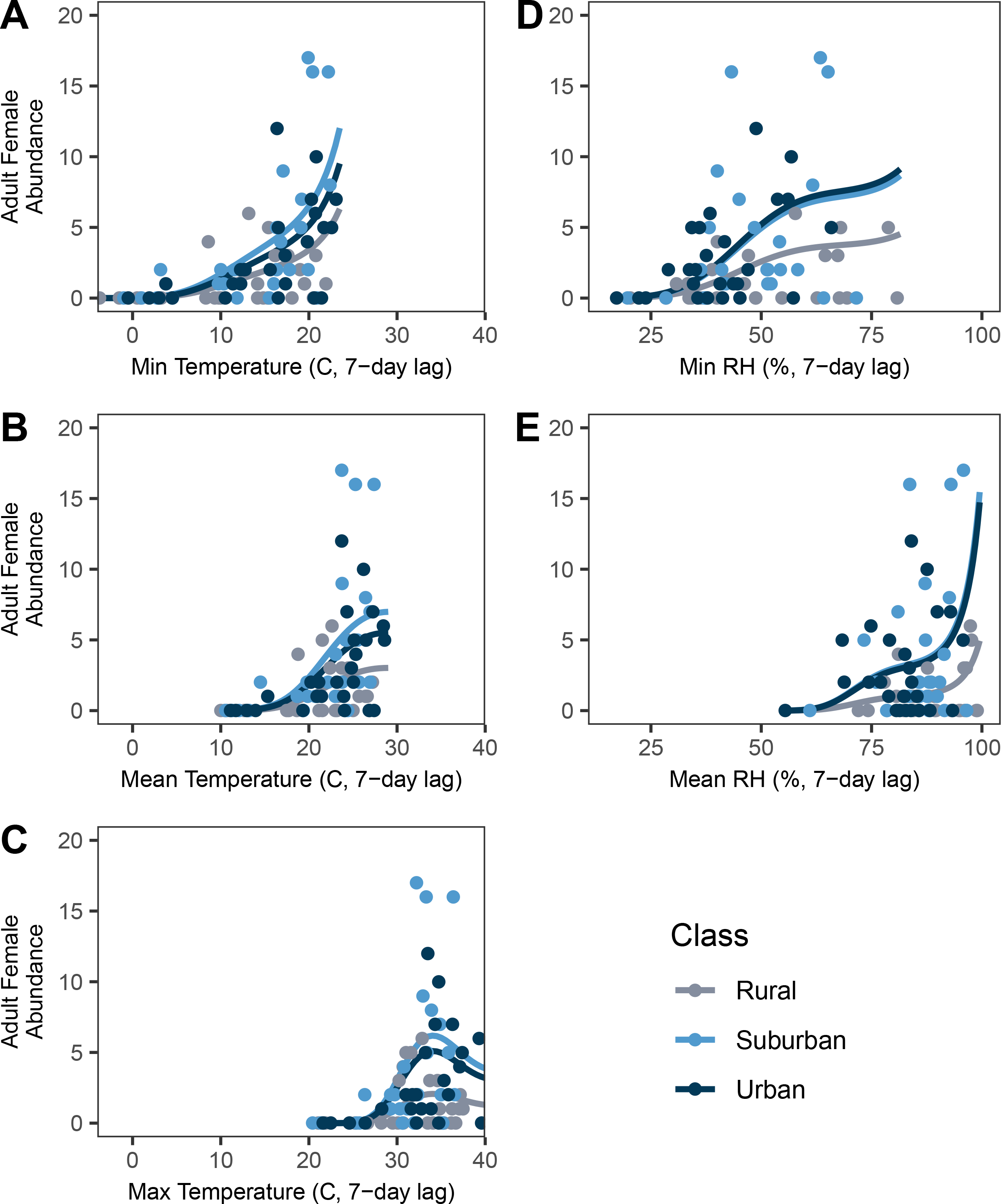
Functional relationship between microclimate variables and adult female abundance for a representative site of each land class. Functional relationships were the same across all land classes, and female adult abundance did not differ across land class. Daily minimum, mean, and maximum temperatures are the on the left side (a,b,c) and daily minimum and mean relative humidity are on the right side (d,e). Lines represent fitted regression lines and raw data are represented by the circles. Because maximum relative humidity did not vary, no regression line was fitted. The suburban and urban fitted curves for minimum relative humidity are visually indistinguishable, and so the suburban curve has been shifted downward for visibility.

### Larval Habitat Density and Adult Abundance

We detected a significant positive relationship between larval habitat density and adult abundance at a site (χ^2^ = 17.788, *df* = 1, p<0.0001), although the effect size was highly dependent on site. Site-level random effects ranged from −0.975 to 1.280, compared to a regression coefficient of 0.364, suggesting that unmeasured covariates at the site level are also contributing substantially to adult abundances. This is further supported by the relatively low model fit (R^2^ = 0.39) compared to the univariate models of microclimate variables described above.

## DISCUSSION

Spatial predictions of mosquito abundances often rely on temperature-dependent mechanistic models derived from mosquitoes’ thermal performance curves.^17–19^ However, availability of larval habitat can also be a strong determinant of adult mosquito abundances, and few models include these in their predictions (but see ^19^). We found that while both climate and larval habitat influenced adult mosquito abundance, climate was a stronger predictor of adult abundance and the functional relationship between microclimate and adult abundance matches predictions based on theories of ectotherm physiology.^41^ Further, neither adult abundance nor the density of *Ae. albopictus* positive larval habitat varied by urban land class, suggesting that, at least for smaller, residential cities, this variation is not significant, and models do not need to differentiate across land class in urban areas.

While we failed to detect a difference in adult abundance across land class, other studies have found mosquito abundances to differ across an urban gradient. Multiple studies that have defined urban gradients according to variation in vegetation density,^42,43^ impervious surface,^44^ or land cover classifications ^45–48^ have found mosquito abundances to vary across these gradients. Li *et al.* focused specifically on *Ae. albopictus* and found adult abundance to increase with increasing urbanization, driven by an increase in larval habitat density.^26^ Our study, however, found no evidence for a difference in positive larval habitat density across land classes, which may explain why we failed to detect a difference in adult abundance. A semi-field experiment conducted at the same study sites as this experiment estimated lower *Ae. albopictus* per-capita growth rates on urban sites compared to rural and suburban sites, driven by lower larval survival rates and smaller wing lengths of emerged adults (a predictor of fecundity) on urban sites.^17^ Taken together, these findings suggest that per capita growth rates may not scale-up to site level population abundances. Other factors, such as the quantity and quality of larval habitat or the availability of hosts for blood feeding,^49^ may further mediate the relationship between container-level growth rates and site level abundances.

Temperature and relative humidity are likely the key variables driving seasonal trends in mosquito density, as they were important predictors of both the density of larval habitats positive for *Ae. albopictus* larvae and adult *Ae. albopictus* abundance. While many studies have observed seasonality in larval habitat abundance,^50,51^ few have directly paired these data with climate variables. The density of larval habitat had an exponentially increasing relationship with temperature, and, indeed, larval habitat was most abundant during the summer sampling periods. Hotter temperatures can increase mosquito biting rates and shorten gonotrophic cycles,^14^ potentially leading to higher oviposition rates and a higher density of larval habitats. Relative humidity, in comparison, had a unimodal relationship with larval habitat, with the number of larval habitats decreasing at high minimum and mean relative humidity. Very few studies have investigated the effects of relative humidity on larval mosquito dynamics. However, Murdock *et al.* found that increases in relative humidity reduced larval survival through a suggested decrease in the surface tension of aquatic environments.^27^ Another explanation is that high relative humidity is associated with strong rainfall events,^52^ which can lower the density of positive larval habitat through flushing events and oviposition avoidance.^53^ Interestingly, these functional relationships (Fig. 4) differed qualitatively from those between climate and adult abundance (Fig. 5), suggesting the effects of temperature and relative humidity on mosquito populations may differ across life stages (ovipositing and hatching vs. emergence and adult survival).

Temperature and relative humidity were also key predictors of adult abundance. The non-linear functions used in the temperature models match the unimodal functional form between ectotherm growth and temperature expected from physiological theory and empirical work in mosquito systems.^17,41^ *Ae. albopictus* abundance was zero at mean temperatures below 10°C, and increased to a peak temperature around 25°C. This agrees with other studies in urban areas that found the minimum threshold for adult activity to be 10°C and laboratory predictions of the optimum temperature of 25°C.^54,55^ In the case of the daily maximum temperature, temperatures during our study period exceeded the optimal temperature for *Ae. albopictus*, and adult abundance decreased at high temperatures, creating a hump-shaped curve. These field findings match general expectations of thermal performance curves derived from laboratory experiments, suggesting that empirically-derived thermal performance curves are applicable to mosquito populations in field settings. Adult abundance also increased with increasing levels of relative humidity. Another study observed adult *Ae. albopictus* mortality rates to decrease with increasing humidity in the field.^56^ This relationship between adult mortality and humidity may drive the positive relationship between relative humidity and adult abundance in our study. We found a positive, though weak, relationship between the density of positive larval habitat and adult mosquito densities. This is in agreement with other studies that have found larval habitat to be predictive of adult densities.^26,31^ In addition to providing more space and resources for immature mosquitoes, high densities of larval habitat can also reduce the time spent searching for oviposition sites, shortening gonotrophic cycles and increasing population growth rates.^57^ The overall performance of the model including larval habitat was lower than one based solely on microclimate. This implies that, while adult abundance and larval habitat are correlated, microclimate alone may more accurately predict mosquito abundances.

By spanning two years, we replicated seasonality, but only across a limited number of sites. While we classified sites into urban land classes based on determinants of microclimate, namely impervious surface, unmeasured site-level characteristics were an important driver of *Ae. albopictus* abundance. For example, one urban site produced more than two-fold the number mosquitoes of any other urban site. This site received daily irrigation throughout the summer months, perhaps contributing to high *Ae. albopictus* abundances, as has been found in *Culex spp*.^58^ The types of artificial containers can differ across socio-economic levels in urban areas.^42^ The type of larval habitat in our study varied widely across sites, from natural bodies such as ponds and treeholes to artificial containers such as flower pots and tires, but there was no pattern across land class (Table S1). Suburban and urban sites in particular had wide variation in habitat types, and the inclusion of social variables such as parcel value or income in our classification could lead to higher uniformity in land classifications.

We found that adult abundance was well predicted by microclimate variables and that the functional relationship between temperature and adult abundance matched that proposed by theory and empirical studies. This study contributes to a small number of studies exploring predictors of *Ae. albopictus* abundance in cities.^26,42^ Unlike past studies, we found no evidence for an effect of urban land class on *Ae. albopictus* abundances, suggesting that city-scale predictive models may not need to explicitly incorporate differences across land classes. However, Athens, GA is a small city, with an average impervious surface of 10% and a population of 127,064, and these results may not apply to larger cities with wider variation in land class, which can differ in temperature by over 5°C.^59^ Future work could expand field studies to additional cities to test the generalizability of these findings and identify contexts (e.g. tropical vs. temperate cities, small vs. large cities) in which these results differ. By pairing mosquito surveys with the collection of microclimate data, our findings support the continued use of temperature-dependent mechanistic models in the spatial prediction of mosquito abundances and mosquito-borne disease risk.

## ACKNOWLEDGEMENTS

We thank members of the Murdock lab for their discussion and field support conducting this study, particularly Diana Diaz. We also thank the property owners who provided us permission to enter their properties to conduct surveys.

## AUTHOR ADDRESSES

Michelle V. Evans: University of Georgia, Athens, GA, USA

Carl W. Hintz: North Carolina State University, Raleigh, NC, USA

Lindsey Jones: Albany State University, Albany, GA, USA

Justine Shiau: University of Georgia, Athens, GA, USA

Nicole Solano: University of Georgia, Athens, GA, USA

John M. Drake: University of Georgia, Athens, GA, USA

Courtney C. Murdock: University of Georgia, Athens, GA, USA

