## Supplemental Table 1 for "Microclimate and larval habitat density predict adult *Aedes albopictus* abundance in urban areas"

**SUPPLEMENTAL TABLES**

|  | **Artificial Container** | **Ground Pool** | **Pond** | **Rockpool** | **Treehole** |
| --- | --- | --- | --- | --- | --- |
| **Rural** | 46/201 | 4/12 | 1/6 | 0/0 | 0/5 |
| **Suburban** | 111/378 | 0/5 | 0/20 | 1/2 | 3/32 |
| **Urban** | 28/129 | 0/22 | 1/35 | 0/0 | 17/38 |

**Table S1.** Distribution of container types across land classes (positive/total). Most container types were found across all land classes, with the exception of rockpools.
